# Multi-color 4D superresolution light-sheet microscopy reveals organelle interactions at isotropic 100-nm resolution and sub-second timescales

**DOI:** 10.1101/2021.05.09.443230

**Authors:** Yuxuan Zhao, Meng Zhang, Wenting Zhang, Qing Liu, Peng Wang, Rong Chen, Peng Fei, Yu-Hui Zhang

**Author notes:** These authors contributed equally: Yuxuan Zhao, Meng Zhang, Wenting Zhang. Correspondence and requests for materials should be addressed to Y.-H.Z. or to P.F.

## Abstract

Long-term visualization of the dynamic organelle-organelle or protein-organelle interactions throughout the three-dimensional space of whole live cells is essential to better understand their functions, but this task remains challenging due to the limitations of existing three-dimensional fluorescence microscopy techniques, such as an insufficient axial resolution, low volumetric imaging rate, and photobleaching. Here, we present the combination of a progressive deep-learning superresolution strategy with a dual-ring-modulated SPIM design capable of visualizing the dynamics of intracellular organelles in live cells for hours at an isotropic spatial resolution of ∼100 nm in three dimensions and a temporal resolution up to ∼17 Hz. With a compelling spatiotemporal resolution, we substantially reveal the complex spatial relationships and interactions between the endoplasmic reticulum (ER) and mitochondria throughout live cells, providing new insights into ER-mediated mitochondrial division. We also localized the motion of Drp1 oligomers in three dimensions and observed Drp1-mediated mitochondrial branching for the first time.

## Introduction

The interactions between diverse organelles or their interactions with intracellular proteins play important roles in many crucial biological processes. A thorough understanding of these interactions is essential for fundamental research in biology and medicine^1-3^. These interactions (i.e., those involving direct physical contact rather than signaling pathways) occur at nanoscale membrane contact sites and rapidly change over a long term throughout the three-dimensional (3D) space of whole cells^4^. Therefore, precise visualization of these interactions not only requires multicolor live cell imaging at a sufficiently high spatial resolution in all three dimensions and a z-depth closer to the full thickness of whole cells (∼5 µm) but also requires long-duration imaging with a low phototoxicity at a high speed to capture the entire process of these rapid dynamic changes^4-7^.

Recently developed 3D superresolution methods have achieved volumetric nanoscale fluorescence imaging and opened up new avenues to address subcellular events previously indiscernible^8-18^. A series of 3D imaging methods based on the well-known single-molecule localization microscopy (3D SMLM) technique have attained extremely high spatial resolutions (up to 10 nm) but at the expense of the temporal resolution with a notable phototoxicity, making these methods difficult to apply in live cell dynamic imaging^8,9^. Compared to 3D SMLM, 3D stimulated emission depletion microscopy (3D STED)^12^ and its latest modification using parallelized reversible saturable optical linear fluorescence transition (3D pRESOLFT), achieve a significantly improved temporal resolution, albeit still only a minute-scale resolution^13^. 3D wide-field structured illumination microscopy (3D SIM) obtains 15 images in each plane, thus doubling the spatial resolution^10,11^. However, the wide-field illumination and multiframe acquisition mode notably increase photodamage and reduce the imaging speed, which seriously limit its applications in long-term time-lapse imaging of whole-cell dynamics^5-7^.

Light-sheet fluorescence microscopy (LSFM) has recently emerged as a technique-of-choice for the fast, 3D imaging of live cells^14,15,19-21^. Among the various techniques, lattice light-sheet microscopy (LLSM) has become the best-known modality owing to its high axial resolution and low phototoxicity^14^. Combined with 3D structure illumination, the spatial resolution of LLSM (LLSM-SIM) can be further improved by a factor of approximately 1.5, at the expense of 5 repetitive acquisitions under a phase-shifted lattice pattern^14^. As a reference, the typical performance of LLSM-SIM is 7.5 s per single cell at a 150×230×280-nm^3^ resolution, and hundreds of volumes are recorded during a period of tens of minutes^14^. Combining LLSM with patterned activation nonlinear-SIM (PA LLSM-SIM) could further improve the spatial resolution to 118×230×170-nm^3^ but at the cost of the temporal resolution (approximately 30-70 s/single cell)^15^. Although most cutting-edge LSFM applications as such successfully reveal organelle interactions, the spatiotemporal performance remains unsatisfactory, given that many 3D intracellular events occur on subsecond timescales.

To further push the limits of superresolution microscopy techniques for the rapid, sustained 3D imaging of live cells beyond the diffraction limit, two major challenges must be addressed^22^: one challenge involves the development of plane illumination towards perfect conditions, which could be achieved through the ready generation of ultrathin chromatic laser sheets without excessive side-lobe excitation and within a sufficiently long range of single cells; the other challenge is superresolution realization with very few diffraction-limited measurements to increase the temporal resolution and reduce phototoxicity. Recently, several types of nondiffraction plane illumination techniques have been developed simply with ring-shape masks or field synthesis^20,21^; however, the lack of advantages in either thinning of the main lobe or suppression of the side lobes over previous LLSM techniques prevents them from further pushing the limit of the axial resolution. Meanwhile, emerging deep-learning techniques are currently involved in various types of 3D microscopy, and they have recently achieved high-quality image restoration based on single-image inputs^23-28^.

Herein, we present an approach that addresses the abovementioned challenges in long-term 3D superresolution imaging of live cells. We developed a 0.45-µm laser sheet illumination approach with low side lobes by designing a dual ring (DR) modulation plate and combined it with an isotropic dual-stage (ID) convolutional neural network to establish the IDDR-selective plane illumination microscopy (IDDR-SPIM) technique that yields superresolution isotropic images based on a single diffraction-limited volumetric input. We also visualized the dynamics of intracellular organelles in live cells for hours (up to 5200 time points, yielding 265200 superresolution images) at an isotropic spatial resolution of ∼100 nm in all three dimensions and a volumetric imaging rate up to 17 Hz. With a compelling spatiotemporal resolution, we successfully captured the morphological changes and locomotion of mitochondria and the endoplasmic reticulum (ER), which might be misinterpreted or remain unresolvable with two-dimensional (2D) superresolution imaging or 3D superresolution imaging at a low speed. Our method substantially revealed the complex spatial relationships and interactions between mitochondria and the ER throughout entire live cells, providing new insights into ER-mediated mitochondrial division. Furthermore, we localized the motion of Drp1 oligomers in three dimensions and observed Drp1-mediated mitochondrial branching for the first time, extending the applications of IDDR-SPIM to a broad field of protein-organelle interactions in cell biology.

## Results

### IDDR-SPIM imaging with a superior spatiotemporal resolution

In contrast to the creation of a thin and uniform illumination plane by dithering a lattice light sheet in the objective space^14^ or scanning a lattice pattern in Fourier space^20^, we generated a scan-free uniform Bessel-type light sheet (i.e., a DR Bessel light sheet) by projecting a line-shaped excitation light onto a delicately designed dual ring modulation plate placed at the back focal plane of the illumination objective (**Supplementary Fig. 1**). By introducing an extra ring and optimizing its inner and outer diameters, we suppressed the side lobes of the generated Bessel light sheet while maintaining a small size of the main lobe. The results reveal that the intensity ratio of the side lobes of a 0.48-µm DR Bessel light sheet reached ∼35%, which is 1.7 times lower than that of the lattice light sheet (**Supplementary Fig. 2**), thus notably reducing any image artifacts generated by strong side lobes (**Supplementary Fig. 3**). Moreover, this simple and cost-effective mask design for the generation of scan-free laser sheets also notably reduced the complexity of the imaging system (**Fig. 1a**). This uniform and low-side-lobe 0.48-µm DR Bessel light sheet attained a light utilization rate of ∼59%, which is ∼1.3 times higher than that of a lattice light sheet of the same thickness (**Supplementary Fig. 2**), and thus achieved an obviously reduced phototoxicity suited for long-term live cell imaging. Furthermore, the DR Bessel light-sheet approach also allowed simultaneous multicolor excitation to image instantaneous organelle-organelle interactions, thereby avoiding the misinterpretations due to sequential dual-color imaging (**Supplementary Fig. 4**).

**Figure 1.**
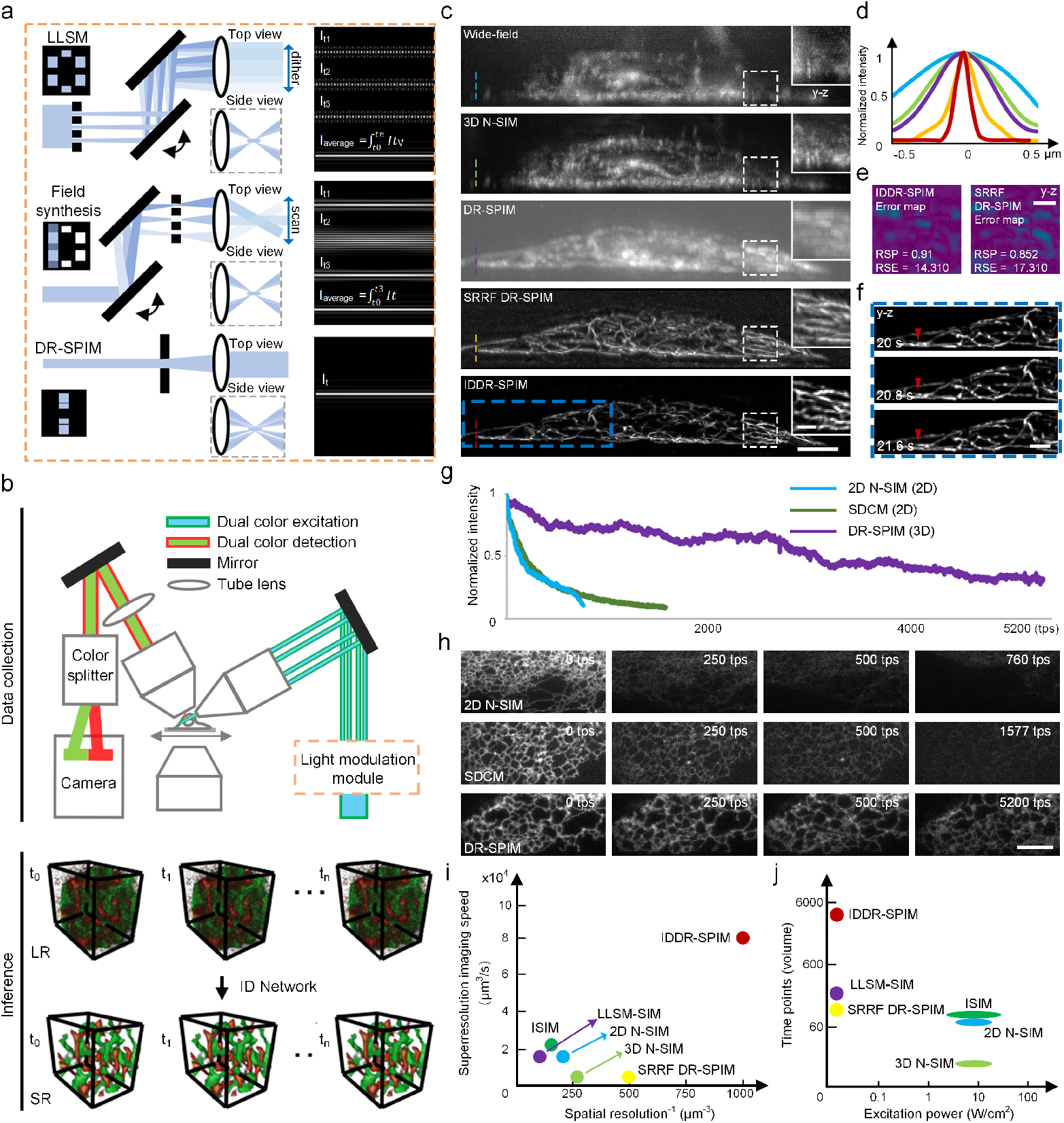
Principle and performance of IDDR-SPIM. **a**, Schematic illustrating the generation of a DR Bessel light sheet and its difference from a lattice light sheet and field synthesis. **b**, Schematic showing the dual-color DR-SPIM setup in conjunction with ID computation for 5D (3D space + time + spectrum) superresolution imaging of intracellular organelles in live cells. LR (diffraction-limited): low resolution, SR: superresolution. **c**, Y-z slices of the microtubules (labeled with Tubulin-Atto 488)^31^ in a live U2OS cell imaged via N-SIM under the wide-field mode (wide-field, numerical aperture (NA)=1.49, top panel) and 3D SIM mode (NA=1.49, 2^nd^ panel), DR-SPIM (3^rd^ panel), SRRF DR-SPIM (4^th^ panel), and IDDR-SPIM (bottom panel). The right white box shows a magnified view of the boxed region for each method. Scale bar, 10 μm and 0.5 μm (insert). **d**, Averages intensity profiles along the lines through 10 single resolved microtubules (shown in the left part of **c**). The blue line, green line, violet line, orange line, and red line indicate the axial resolution of N-SIM under the wide-field mode (NA = 1.49), 3D N-SIM (NA=1.49), DR-SPIM, SRRF DR-SPIM, and IDDR-SPIM, respectively. **e**, Error map and corresponding resolution-scaled error (RSE) and resolution-scaled Pearson coefficient (RSP) of IDDR-SPIM and SRRF DR-SPIM reconstructions compared to the raw DR-SPIM reconstructions. Scale bar, 0.5 μm. **f**, Time-lapse images of the blue box in **c** showing rapidly wiggling microtubules (the red arrows). Scale bar, 1 μm. **g**, Comparison of the photobleaching rates between 2D N-SIM, spinning disk confocal microscopy (SDCM, 2D mode), and IDDR-SPIM. **h**, Corresponding ER images of live U2OS cells imaged at certain time points (tps) via 2D N-SIM (9 frames over 1.8 s at each time point), SDCM (1 frame over 0.2 s at each time point, 2D mode), and IDDR-SPIM (51 frames over 1 s at each time point). Scale bar, 5 μm. **i**, Comparison of the spatiotemporal performance between IDDR-SPIM, SRRF DR-SPIM, LLSM-SIM, 2D N-SIM, 3D-SIM, and ISIM. **j**, Comparison of the applied excitation power and acceptable number of recorded time points between the abovementioned six microscopy methods.

**Figure 2.**
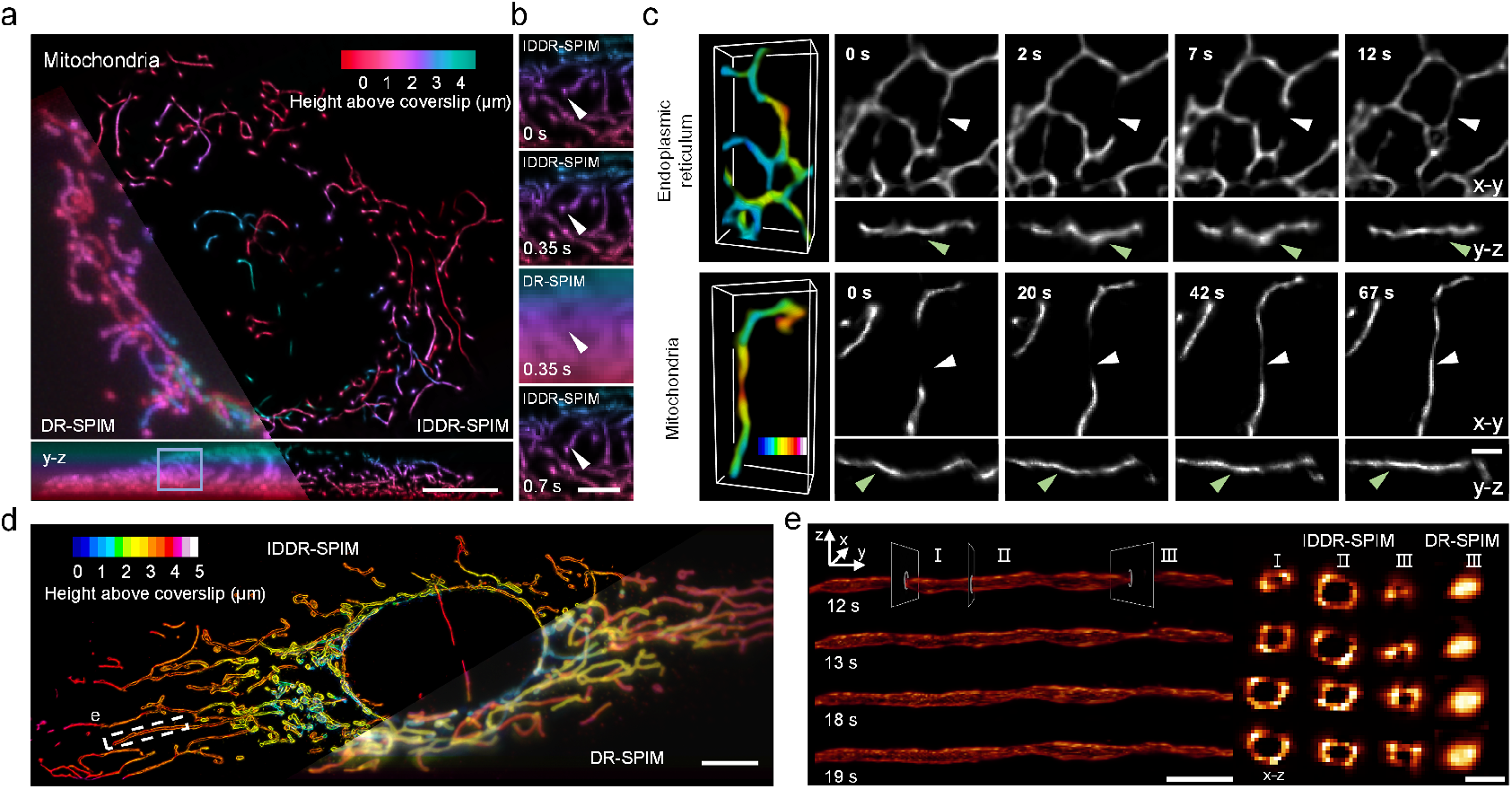
4D superresolution imaging revealing the morphological changes and locomotion of mitochondria or ER in live cells. **a**, Color-coded x-y and y-z MIPs of the mito Ma (tagged with Cox4-mKate2) in an entire live U2OS cell. The left part shows the diffraction-limited DR-SPIM result, and the right part shows the superresolution result enhanced via ID. The whole cell was finally imaged at an ∼3 volumes per second rate and an ∼ 100-nm isotropic resolution (IDDR-SPIM). Scale bar, 10 μm. **b**, Time-lapse magnified views of the blue boxed region in **a**, showing a mitochondrial movement (the white arrows) along the axial plane. These dynamics are well resolved by IDDR-SPIM but are otherwise indistinguishable in the diffraction-limited result (DR-SPIM). Scale bar, 0.5 μm. **c**, 3D visualization (leftmost column) and corresponding x-y and y-z MIPs of the ER (tagged with EGFP-Sec61β) or mito Ma (tagged with Cox4-mKate2), respectively. The ER images in the top row reveal that the axial fluctuation (the green arrows) in the tubular ER might likely be misinterpreted as ER tubule division and reconnection (the white arrows) based on merely 2D imaging. The mitochondrial images in the bottom row reveal that the fluctuation (the green arrows) in the mitochondria might also be misinterpreted as a mitochondrial fusion process (the white arrows). **d**, Color-coded x-y MIPs of the mito OM (tagged with Tomm20-EGFP) in a whole live U2OS cell imaged via diffraction-limited DR-SPIM (right) and IDDR-SPIM (left). The entire cell was imaged at a 1 volume per second rate to capture the dynamics. Scale bar, 10 μm. **e**, Time-lapse 3D visualization of a single mitochondrion retrieved from the boxed region in **d**, showing the morphological changes in the mito OM. The right part shows the x-z sectional planes through the three different sites at the different time points, indicating that in contrast to the diffraction-limited results, which are completely ambiguous (rightmost column), IDDR-SPIM finely resolves the nanometer-scale contraction and expansion dynamics of the mito OM in three dimensions (columns I, II, and III). Scale bar, 2 μm (left) and 0.5 μm (right).

**Figure 3.**
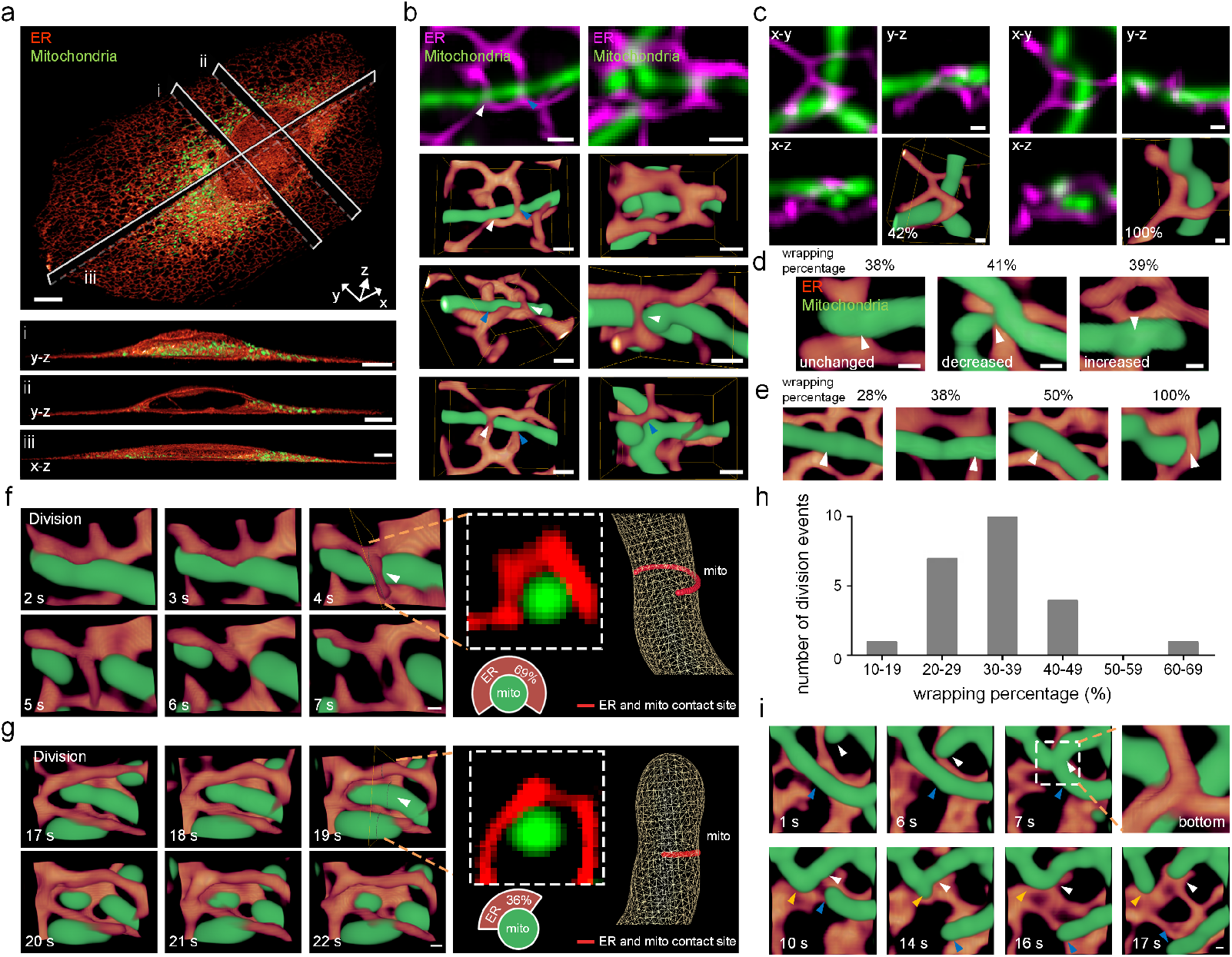
5D superresolution imaging revealing the complex spatial relationships and the dynamic interactions between the ER (red or magenta) and mitochondria (green) in live cells. **a**, Dual-color volume rendering (top panel) of the ER (tagged with EGFP-Sec61β) and mito Ma (tagged with Cox4-mKate2) in a whole live U2OS cell. The lower panels show the y-z and x-z MIPs along the sectional planes in the top panel. Scale bar, 5 μm. **b**, Illustrative ROI, shown as one x-y MIP and three corresponding 3D renderings from different perspectives, revealing the real spatial relationships and contacts between the ER and mitochondria otherwise hardly visible in x-y views (top row). Scale bar, 500 nm. **c**, Illustrative ROI, shown as x-y, y-z, and x-z MIPs and a 3D rendering, revealing the ER wrapping percentages (the white numbers) at the contact sites otherwise indiscernible in the lateral images. Scale bar, 500 nm. **d, e**, Mitochondrial morphology at the ER-mitochondria contact sites with similar or different ER wrapping percentages. Scale bar, 200 nm. The white or blue arrows in **b, d, e** indicate the ER-mitochondria contact sites. **f, g**, Time-lapse dual-color volume renderings showing the dynamic processes of ER-mediated mitochondrial division. The images in the white boxes show the raw reconstructed images of the corresponding y-z sectional planes. The 2D and 3D models show the percentages of the ER wrapping around the mitochondrial circumference (the contact sites marked in red). **h**, Quantitative statistics of the ER wrapping percentage at the division sites during the ER-mediated mitochondrial division events (n=23). **i**, Time-lapse dual-color volume renderings showing the dynamic process involving the cooperation of the ER and one mitochondrion (the white arrows) in the division of another mitochondrion (the blue and yellow arrows). The zoomed-in view shows the contact site where the ER occurs in conjunction with the two mitochondria. Scale bar, 200 nm.

**Figure 4.**
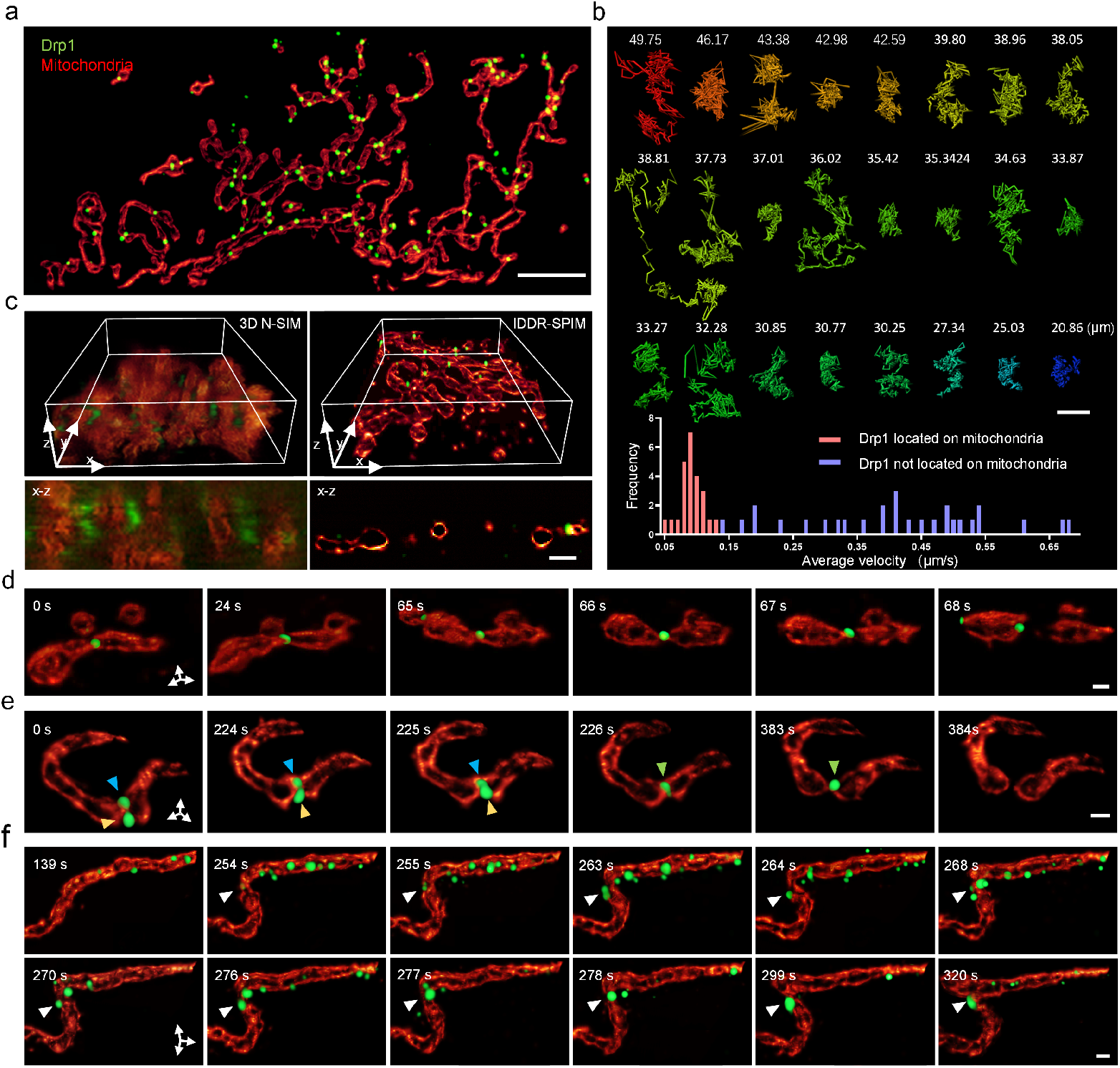
5D superresolution imaging revealing the dynamic interactions between the Drp1 oligomers (green) and mitochondria (red). **a**, A dual-color volume rendering of the Drp1 oligomers (tagged with mCherry) and mito OM (tagged with Tomm20-EGFP) in a live U2OS cell. Scale bar, 5 μm. **b**, Trajectories of the Drp1 oligomers located on the mitochondria, which were randomly selected from the Drp1 oligomers in **a**, and quantitative statistics of the average velocities of the Drp1 oligomers (n=24 for the Drp1 oligomers located on the mitochondria and n=28 for the Drp1 oligomers not located on the mitochondria, the trajectories of which are shown in **Supplementary Fig. 13**). The numbers indicate the length of the corresponding trajectories. Scale bar, 0.5 μm. **c**, Dual-color volume renderings and x-z MIPs of the Drp1 oligomers and mito OM imaged via 3D N-SIM or IDDR-SPIM in a live U2OS cell. Scale bar, 1 μm. **d, e, f**, Time-lapse dual-color volume renderings showing the dynamic processes of Drp1-mediated mitochondrial division (**d, e**) or Drp1-mediated mitochondrial branching (**f**). The arrows indicate the division sites or the branching sites. Scale bar, 200 nm.

With the use of the DR Bessel light sheet, we obtain high-quality 3D images of a whole cell at an ∼230×230×450-nm^3^ native resolution and up to a 340-ms temporal resolution. Further computational superresolution imaging exceeding the diffraction limit then requires the input of multiple stacks, which inevitably increases the acquisition time and phototoxicity^14,15^. To circumvent this dilemma, we combined our dual-stage superresolution model^28^ with content-aware restoration (CARE)^23^ to create an isotropic superresolution neural network, namely, the isotropic dual-stage (ID) neural network, that finely resolved the raw DR Bessel light-sheet image into an ∼100-nm isotropic resolution based on a single diffraction-limited volumetric input. We first obtained 3D superresolution radial fluctuation (3D-SRRF)^29,30^ images of the ER, microtubules, mitochondrial outer membrane (mito OM), and mitochondrial matrix (mito Ma) in fixed cells. These datasets were paired with their corresponding diffraction-limited datasets to train the ID neural network so that the well-established neural network converted a diffraction-limited anisotropic image with a low signal-to-noise ratio (SNR) into an isotropic superresolution image with a high SNR. Via DR Bessel light-sheet imaging and ID computation, we establish the IDDR-SPIM method that achieves 3D video recording of intracellular organelle dynamics with an overwhelming spatiotemporal performance and a high fidelity (**Fig. 1b, Supplementary Fig. 5**). We imaged microtubules within an entire live U2OS cell via the Nikon-SIM (N-SIM) under the wide-field mode (wide-field) or 3D SIM mode (3D N-SIM), DR-SPIM, SRRF DR-SPIM, and IDDR-SPIM and compared the reconstructed y-z planes of the cell (**Fig. 1c**). The wide-field, 3D N-SIM, DR-SPIM, and SRRF DR-SPIM modes attained axial resolutions of 1 μm, 600 nm, 500 nm, and 190 nm, respectively, at volumetric speeds of 4, 60, 0.8, and 24 s/cell, respectively, and the ultimate IDDR-SPIM mode achieved a speed of 0.8 s/cell at an isotropic axial resolution of ∼100 nm (**Fig. 1d**). Moreover, the ID-based reconstructions were sufficiently accurate, with well-validated SQUIRREL^32^ analysis metrics (**Fig. 1e, Supplementary Fig. 6**). The high spatiotemporal resolution therefore exclusively allowed the visualization of single wiggling microtubules at a subsecond rate (**Fig. 1f, Supplementary Video 1**). In addition to an isotropic superresolution, the strong capability of low-SNR data restoration accomplished under a low-intensity DR Bessel light sheet (∼1 W/cm^2^) achieved an ∼5-fold improved SNR in the restored output (**Supplementary Fig. 7**), thus further permitting 3D superresolution imaging of the ER in a live cell at 5200 time points, yielding 265200 superresolution images (51 planes/volume at each time point) with less than 0.01% photobleaching per volume (**Fig. 1g, Supplementary Video 2**). In contrast to the ultralow phototoxicity of IDDR-SPIM imaging, the fluorescence signals acquired by either spinning disk confocal microscopy (SDCM) or 2D N-SIM were obviously bleached after a few hundred time points even though merely 1 or 9 frames, respectively, were acquired at each time point in these two implementations (**Fig. 1g, 1h, Supplementary Video 3**). Then, we compared the spatiotemporal performance of IDDR-SPIM to that of LLSM-SIM^14^, 2D N-SIM, 3D N-SIM, instant SIM (ISIM, the fastest SIM method currently available)^16,17^, and SRRF DR-SPIM (**Fig. 1i**). IDDR-SPIM attained a higher spatial resolution than that of the other state-of-the-art superresolution methods. Moreover, the imaging speed of ISIM is comparable to that of IDDR-SPIM, but its spatial resolution of ∼145×145×350 nm^3^ is much lower. We also noted that the advantages of IDDR-SPIM in regard to low-side-lobe illumination and single-image superresolution (SISR) enabled sustained 3D imaging at thousands of time points, which is at least one order of magnitude higher than that of the compared approaches (**Fig. 1j**).

### 4D superresolution imaging of dynamic organelles in live cells

We performed 4D (3D space + time) superresolution imaging of the dynamics of the ER and mitochondrial networks via IDDR-SPIM. Cox4-mKate2-tagged mitochondrial matrix in whole live U2OS cells were recorded at a volumetric imaging rate of 3 Hz (**Fig. 2a, Supplementary Video 4**). After ID enhancement was applied, we clearly identified the axial locomotion of single mitochondria, which could hardly be discerned via diffraction-limited imaging (**Fig. 2b**). By reducing the size and number of frames acquired at each time point, we further captured the dynamic processes of the ER tubules (tagged with EGFP-Sec61β) rapidly moving in a specific region of interest (ROI, ∼10×200×12.8 μm^3^) at a video rate of ∼17 Hz, thus successfully revealing nanometer-scale 3D oscillations previously unresolvable with other imaging approaches **(Supplementary Fig. 8, Supplementary Video 5)**. In addition to the high spatial and temporal resolutions, true 3D imaging is crucial to correctly interpret the locomotion of organelles, especially when they move along the axial direction. As shown in **Fig. 2c**, we projected 1-μm depth (depth of focus (DOF) of a 1.49 NA objective) IDDR-SPIM image stacks of the ER or mitochondria to simulate the effective signals acquired via 2D SRRF microscopy (x-y maximum-intensity projections (MIPs)). With merely 2D superresolution images provided, the axial movements of the ER and mitochondria would likely be misinterpreted as division or fusion (the white arrows in the x-y MIPs). The third dimension revealed by IDDR-SPIM herein is necessary to accurately identify the dynamics (the green arrows in the y-z planes). We further imaged the mito OM in a whole live cell (tagged with Tomm20-EGFP) (135 planes/s) (**Fig. 2d**). The dynamics of the mitochondrial networks were *in toto* visualized (**Supplementary Video 6**) with the fine hollow structures of a single mitochondrion (minimum diameter less than 100 nm), which were ambiguous in the raw DR-SPIM results, and they were three-dimensionally superresolved via IDDR-SPIM (**Fig. 2e**). Therefore, we clearly observed the process of morphological change occurring in a single mitochondrion, such as contraction and expansion, along the three dissected sectional planes (I, II, and III in **Fig. 2e**).

### 5D superresolution imaging quantitatively reveals the interactions between the ER and mitochondria

With the use of IDDR-SPIM, we also achieved simultaneous 5D (3D space + time + spectrum) superresolution imaging of the ER and mitochondria throughout a whole live U2OS cell at a speed of 188 planes/s at 300 time points (56400 dual-color superresolution images, **Fig. 3a, Supplementary Video 7**). The excellent spatial resolutions along all three axes enabled us to properly distinguish the complex spatial relationships between the ER and mitochondria (**Fig. 3b**). While the projection images finely show the ER and mitochondrial structures, it is difficult to interpret the real spatial relationships between them (**Fig. 3b**). The 3D renderings of these regions allowed a clear depiction of the spatial entanglements, highlighting the need for true 3D imaging to better understand the complex interactions between organelles (**Fig. 3b**). With this new precision, we quantified the percentage of the mitochondrial circumference wrapped by the ER (i.e., the ER wrapping percentage) at the contact sites (**Supplementary Fig. 9**), which was not discernible in the lateral images (**Fig. 3c**). We found that at the contact sites with similar ER wrapping percentages (38-41%), the mitochondrial diameter substantially decreased in some subgroups but increased or remained unchanged in others. The mitochondrial diameter also remained unaltered when the ER wrapping percentage notably varied (28-100%, **Fig. 3c-e, Supplementary Fig. 10**). These results suggest that there exists no obvious causal relationship between mitochondrial constriction and ER wrapping.

The compelling ability of our 5D superresolution imaging technique also sheds new light on the dynamic behaviors of organelle interactions. The true 3D views clearly reveal a dynamic process of ER-mediated mitochondrial division, in which the ER tubule extended and wrapped around the mitochondrion, after which the mitochondrion divided within the next 1 s, followed by the retraction of the ER tubule (**Fig. 3f, Supplementary Video 8**). Moreover, the ER wrapping percentages at the division sites differed, ranging from 18% to 69%, mostly lower than 50% (n =23) (**Fig. 3f-h, Supplementary Video 9**). Previous studies have shown that ER-mitochondrial contacts play active roles in mitochondrial division^33,34^. Two hypotheses have been proposed in terms of the regulation of mitochondrial division by ER tubules^33-35^: (I) The physical wrapping of ER tubules around mitochondria might exert a localized mechanical force to constrict the mitochondria before the division machinery is recruited; (II) the ER does not cause mitochondrial constriction but simply interacts with the mitochondria and provides biochemical signals and components. Our results (**Fig. 3d, Fig. 3f-h**) suggest that there occurs no obvious causal relationship between mitochondrial contraction and ER wrapping and that the ER wrapping percentages at the sites of mitochondrial division greatly vary, suggesting that ER-mitochondrial contacts might function as a platform for the recruitment of the mitochondrial division machinery (hypothesis II) rather than physical mitochondrial constriction (hypothesis I).

We also observed a dynamic process in which one mitochondrion came into contact with the ER and another mitochondrion, followed by the subsequent division of the mitochondrion and a subsequent quick exit of the other mitochondrion, indicating that mitochondria may also cooperate with the ER to trigger the division of another mitochondrion (**Fig. 3i, supplementary video 10**). To our knowledge, to date, there have been no reports that mitochondria themselves might also cooperate with the ER in other mitochondrial division processes. Therefore, our results might provide a new perspective on ER-mitochondrial interactions. Considered together, the observations of these distinct dynamic processes demonstrate the capability of IDDR-SPIM to reveal the complex spatial relationships and interactions between organelles at a novel 5D level.

### 5D superresolution imaging quantitatively reveals the interactions between Drp1 oligomers and mitochondria

To further extend our technique to studies of protein-organelle interactions, we tested the performance of IDDR-SPIM in 5D superresolution imaging of Drp1 oligomers (oligomers of Drp1 proteins tagged with mCherry) and mitochondria (tagged with Tomm20-EGFP) in live cells (**Fig. 4a, Supplementary Fig. 11**). Compared to organelles, protein oligomers usually possess smaller sizes and therefore contain fewer tagged fluorescent proteins, which results in faster photobleaching, thus making long-term visualization of protein-organelle interactions generally more difficult than that of organelle-organelle interactions.

Owing to the high spatiotemporal resolution and low phototoxicity of IDDR-SPIM, we obtained 5D superresolution images of the Drp1 oligomers and mitochondria at up to 400 time points and a volumetric imaging rate of 1 Hz (79 planes recorded over 1 s at each time point). The rapid movements of the Drp1 oligomers along all three spatial directions were *in toto* visualized with tracking times and lengths of the trajectories up to 400 s and 49.75 µm, respectively (**Fig. 4b**), which were very blurry under spinning disk confocal microscopy (**Supplementary Fig. 12)** or conventional superresolution microscopy (e.g., 3D N-SIM, **Fig. 4c**). We also clearly resolved the spatial relationship between the Drp1 oligomers and mitochondria, and quantitative analysis indicated that the average velocities of the Drp1 oligomers not located on the mitochondria were much higher than those of the Drp1 oligomers located on the mitochondria, and the range of variation was also larger (**Fig. 4b, Supplementary Fig. 13**).

The 3D details of the fine mitochondrial hollow structures and individual Drp1 oligomers during Drp1-mediated mitochondrial division were also explicitly visualized and traceable (**Fig. 4d, 4e**). Two Drp1 oligomer puncta were observed to constrict the outer membrane of the mitochondrion, which then fused, and mitochondrial division eventually occurred (**Fig. 4e, Supplementary Video 11**), thus providing spatiotemporal details of the regulation effect of Drp1 oligomers on mitochondrial division. Notably, we recorded the process of Drp1-mediated mitochondrial branching (**Fig. 4f, Supplementary Video 12**). The Drp1 oligomers located on a mitochondrion translocated along it, accumulated at the branching site, and then mediated mitochondrial constriction, followed by branching rather than division. Drp1 oligomers have been reported to play a key role in mitochondrial division^33^, but to our knowledge, this is the first time that Drp1 was observed to play an important role in mitochondrial branching. These results demonstrate the ability of IDDR-SPIM to capture and discern protein-organelle interactions, verifying that this approach is of great importance to cell biology research.

## Discussion

In summary, we developed a DR Bessel light-sheet system, which could easily generate thin (0.45-nm), multicolor (488 and 556 nm) light sheets with relatively low side lobes within a sufficiently long illumination range of the entire cell. The DR Bessel light-sheet design is free from scanning common in field synthesis or from dithering common in LLSM, thus eliminating the use of precise scanning equipment. Additionally, compared to the previous single ring Bessel light-sheet design^21^, the addition of the second ring was validated to efficiently reduce the side lobes. The DR Bessel light-sheet system allowed simultaneous dual-color 3D imaging at a high axial resolution verified by the raw data. In addition to the new hardware design, we also developed a comprehensive deep-learning mode to computationally surpass the diffraction limit and improve the contrast based on a single diffraction-limited and noisy input, thereby achieving an ∼100-nm isotropic superresolution and an ∼5-fold improved SNR in the restored output. Furthermore, the high utilization rate of the excitation light and the notable capability of the ID neural network for the restoration of low-SNR data acquired under low-intensity illumination together yield an ultrahigh photon efficiency, which thus leads to a notably reduced phototoxicity well suited for long-term live cell imaging over thousands of cycles. Therefore, the combination of the DR Bessel light-sheet illumination with ID neural network restoration enabled the proposed IDDR-SPIM technique to achieve sustained volumetric imaging of intracellular organelles and proteins interacting in whole live cells at an ∼100-nm isotropic resolution along all three axes, at up to a 3-Hz volumetric imaging rate, and with less than 0.01% photobleaching per 3D measurement, together achieving an overall spatiotemporal performance far superior to that of current leading 3D superresolution microscopy techniques (**Supplementary Table 1**). We envision that through further combination with an axial scanning scheme^36^ or the nonlinear optical effect^19^, an even smaller thickness of the light sheet and complete elimination of side-lobe excitation could be further accomplished to allow LSFM imaging towards perfect illumination conditions. We also note that while ID restoration improved the reconstruction quality from the diffraction-limited resolution determined by the optical system to the superresolution obtained via the training procedure, this required the preacquisition of a considerable number of superresolution images for training. We expect this will be improved with the continued development of deep-learning techniques, which aim to accomplish a strong generalization ability and weak training supervision, thus allowing model implementation with much fewer required training data.

The compelling spatiotemporal resolution and the simultaneous dual-color imaging ability of IDDR-SPIM allow the proper distinction of the complex spatial relationships and dynamic interactions between the ER and mitochondria, thus providing a new perspective on ER-mediated mitochondrial division and demonstrating its excellent capability in revealing dynamic organelle-organelle complex interactions. The successful recording of the dynamic interactions between Drp1 oligomers and mitochondria further extends the applications of the IDDR-SPIM method to a broad field of protein-organelle interactions in cell biology. We believe that this technique will be of great use in a series of new functional and structural studies of cell biology aiming to analyze the 3D distribution and dynamic interactions between organelles and proteins in intact live cells. Here, we only demonstrated simultaneous dual-color imaging of live cells, and we anticipate that in the future, more channels could be combined in the same manner to assist in studying more complex interactions among organelles and intracellular molecules (e.g., three or four).

## Online content

Any methods, additional references, and supplementary information are available on line.

## Acknowledgments

We thank the Optical Bioimaging Core Facility of WNLO-HUST for the support in data acquisition, and the Analytical and Testing Center of HUST for spectral measurements. This work was supported by the following grants: National Natural Science Foundation of China (Grant nos. 92054110, 61827825, 21874052, 21927802, and 31770924), National Key R&D program of China (Grant no. 2017YFA0700501), Science Fund for Creative Research Group of China (Grant No. 61721092), Innovation Fund of WNLO, the Fundamental Research Funds for the Central Universities (Grant no. 2018JYCXJJ021), the Junior Thousand Talents Program of China (P. F.), and Director Fund of WNLO.

## Author contributions

Y.-H.Z. and P.F. conceived and oversaw the project. Y.Z., Q.L. and R.C. developed the optical setups. Y.Z. and Q.L. developed the programs. M.Z. and W.Z. prepared the experimental samples. Y.Z., M.Z., W. Z., Q.L., and P. W. processed the images and analyzed the data. Y.Z., M.Z., P.F. and Y.-H.Z. wrote the manuscript with discussion and improvements from all authors.

## Competing interests

The authors declare no competing interests.

## Methods

### Dual ring Bessel light sheet microscopy

The microscope was built on an optical table as illustrated in **Supplementary Fig. 1**. Two laser beams of 488 nm and 556 nm from single-mode fiber were collimated by the objective (PLanFLN×10/0.3, Nikon). After that, the laser was passed through three pairs of cylindrical lenses with a focal length ratio of 1: 5 (f_1a_= 50 mm, f_2a_ = 250 mm and f_1b_ = 50 mm, f_2b_ = 150 mm, f _3b_= 150 mm, f_4b_ = 250 mm, Thorlabs) to expand the beam diameter around 5 times. The expanded beam was further focused by a cylindrical lens (f = 250, Thorlabs). And the customized dual-ring optical mask was placed at the focus of the cylindrical lens to provide spatial filtration of the line-shape beam. An achromatic lens with 75-mm focal lens and an infinity-corrected objective (TTL100, Thorlabs) was used to relay the optical pattern to the back focal plane of illumination objective. The optical pattern was finally transformed by the excitation objective to create the desired Bessel light sheet at its front focal plane. The fluorescence signals generated within the specimen were collected by a detection objective (LUMFLN×60/1.1W, Olympus), the focal plane of which was the plane of light-sheet illumination. A 200mm tube lens (ITL200, Thorlabs) was used to provide an overall magnification of 66.7X. We used an image splitter (W-VIEW GEMINI, HAMAMATSU) to simultaneously collect two-wavelength images with one sCMOS camera (Orca Flash4.0 v3, Hamamatsu). The sample motion part, include the high-speed sample piezo (P621.CD, Physical instrument), a trio of closed-loop micro-positioning stages (L-505, U-521.24, physical instrument) and the epifluorescence detection optical path (LUMPLanFLN×20/0.45, Olympus and TTL180, Olympus) are similar with lattice microscopy setup^14^. During image acquisition, the sample stage was activated in a stepwise mode and the camera acquired one image till the stage was stabilized at a desired position. The camera exposure was triggered with an ∼ 2-ms delay to further avoid the motion blur, and the laser was turned off when the sample was in the middle of movement. The analog control signals sent to the camera and the piezo stage were generated by a data acquisition device (PCIe-6259, National Instruments).

### Training data acquisition and isotropic dual-stage (ID) convolutional neural network validation

ID procedure included our dual-stage 3D superresolution network^28^, integrated with an isotropic axial improvement using CARE model^23^. We first used the DR-SPIM with 450-nm light sheet thickness to consecutively collect 30 groups of high-SNR diffraction-limited image stacks of microtubule, mitochondria matrix, mitochondria outer membrane, and endoplasmic reticulum, on at least 10 individual fixed cells, to generate high-SNR SRRF ground truths for data training on the dual-stage network. After each time of abovementioned repetitive collection, low-SNR diffraction-limited image stacks corresponded to the SRRF results were also acquired with low-intensity illumination. Besides, synthetic high-resolution ground truths and low-resolution 3D images of microbeads were generated to simulate the point-like signals of labelled Drp1 protein. For the dual-stage training of all the organelles / Drp1 protein, we used the down-sampled images of the ground truths as the middle-resolution buffer data, which exhibited more details and higher SNR as compared to the low-resolution datasets. The paired low-resolution, middle-resolution and high-resolution datasets were together used as the training datasets for the training on dual-stage network. For the training data preparation of isotropic enhancement networks, the lateral slices of SRRF images with better resolution were used as high-resolution ground truths and the corresponding low-resolution images was generated by degrading these lateral slices to simulate the anisotropic axial slices.

### Assessment of the resolution improvement and fidelity of isotropic dual-stage (ID) convolutional neural network

We used the resolution-scaled error (RSE) and the resolution-scaled Pearson coefficient (RSP) to assess the three-dimensional fidelity of ID convolutional neural network outputs and SRRF results in both the x-y and y-z planes by using the NanoJ-SQUIRREL^32^. The RSE and RSP were calculated by setting the max intensity projection of raw diffraction-limited DR-SPIM images as the references to be compared with ID and SRRF results. The line plot profiles in x-y and y-z plane of each method were measured by using the Fiji^37^. Then curve fitting in MATLAB was used to calculate the resolution of each methods.

### Simulated property of the DR plane illumination using MATLAB

We simulated the property of DR light sheet from the back focal plane of illumination objective. We assumed the input light wave was *U*_0_and the optical filter at the back focal plane was *F*_*mask*_. Then the optical pattern at the back focal plane is expressed as:

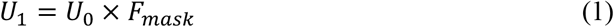

From the Fresnel diffraction integral^38^, the light wave complex amplitude arrives at the objective with propagation distance *f* is:

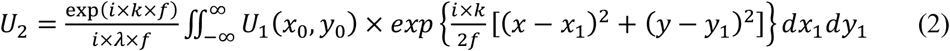

where *f* is focal length of objective lens, *λ* is the wavelength of input light wave. The light wave complex amplitude after the objective lens is:

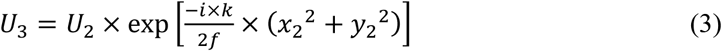

Again, using the Fresnel diffraction integral^38^, the light wave complex amplitude at the focal plane of objective lens is:

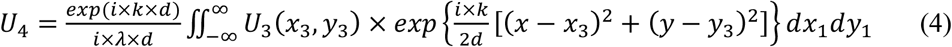

where *d* is the distance away from the objective lens.

To simulate the light sheet property in propagation direction, equation (4) was repeated for different distance *d* from the objective.

### Bessel Light sheet optical parameter optimization

In order to obtained a better optical parameter of low-sidelobe DR Bessel sheet, we fixed the radius and width of outer ring and change the inner ring to optimize the light sheet property. The corresponding simulation program was written in MATLAB.

### Imaging conditions for live-cell experiments

Long term imaging of the endoplasmic reticulum (ER) was implemented, to compare the photon bleaching rate between DR-SPIM, spinning disk confocal microscope and N-SIM microscope. In DR-SPIM experiments, the laser intensity was adjusted to ensure that the image quality was enough for ID reconstruction with high fidelity. In spinning disk microscopy experiments, the laser intensity was carefully adjusted to make the SNR of acquired images similar to the that of DR-SPIM. In Nikon N-SIM experiments, the laser intensity was adjusted to ensure that the images quality was sufficient for superresolution reconstruction. For testing the phototoxicity of DR-SPIM, over 5200 3D image stacks of endoplasmic reticulum (51 slices in each stack, 1 volume per second) were sequentially acquired (The long-term imaging was separated into two stages owing to the computer memory limitation). For spinning disk microscopy, totally 1536 2D slices was acquired consecutively with 100-ms exposure time. For N-SIM in 2D mode, totally 760 2D slices were acquire consecutively with 100ms exposure time.

The bleaching rates were then calculated using Fiji^37^ software. The signal intensity for each method was set as the mean intensities of multiple 50×50 μm areas containing structure signals. The background signals were correspondingly measured in the regions without containing signals. The normalized photon bleaching over time was finally calculated using the following equation:

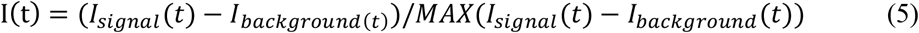

where *I*_*signal*_ and *I*_*background*_ are the mean intensity value of the signal and the background in the image, respectively.

### Quantification of signal to noise ratio (SNR)

The SNR in Supplementary figure 7 was calculated by:

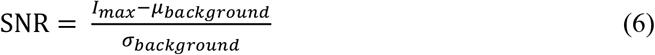

where *I*_*max*_ is the maximum intensity value in the image, *μ*_*background*_ and *σ*_*background*_ are the mean and the standard deviation of the background, respectively^26^.

### N-SIM imaging

N-SIM microscope (Nikon Instruments) was equipped with a 100×/1.49 NA objective and two laser beams (488 nm, 561 nm). The raw images were captured with an EMCCD camera (Andor Technology). For 2D and 3D models, 9 and 15 raw images (three-phase × three-orientation, and five-phase × three-orientation) were acquired to reconstruct the 2D and 3D superresolution images, respectively, using NIS-Elements software. The reconstructed SIM images were analyzed with Fiji^37^ and volumetrically rendered by Amira software.

### The Drp1 oligomers tracing

Trajectories of the Drp1 oligomers were tracked semi-automatically using the commercial Imaris software. The mean speed and the length of the trajectories of the Drp1 oligomers located on the mitochondria were automatic calculated by Imaris after tracking 400 time points at a 1 volume per second rate. For the Drp1 oligomers not located on the mitochondria, the mean speed and the length of the trajectories were calculated by Imaris after tracking at least 50 time points.

### Cell culture and transfection

U2OS cells were grown in culture medium which containing McCoy’s 5A medium (Thermo Fisher Scientific) supplemented with 1% antibiotic-antimycotic (Thermo Fisher Scientific) and 10% fetal bovine serum (FBS; Thermo Fisher Scientific) at 37 °C with 5% CO_2_ in a humidified incubator. The day before transfection, U2OS cells were seeded into the wells of a 24-well plate with 500 μL culture medium. The indicated plasmid was transfected into U2OS cells by the Lipofectamine LTX (Invitrogen) according to the standard protocol. The cells were digested with 0.25% trypsin (Thermo Fisher Scientific) 6-8 hours after transfection, and seeded onto coverslips (customized, Shanghai Jingan Biotechnology Co., Ltd.) and cultured at 37 °C with 5% CO_2_ for another 36 hours. When imaging, cells were cultured in phenol red free McCoy’s 5A medium (customized, Boster Biological Technology co., Ltd.) at 37°C.

### Recombinant plasmids

mCherry2-Sec61β plasmid was generated by replacing EGFP with mCherry2 in the EGFP-Sec61β vector (a gift from Prof. Liangyi Chen (Peking University)). Cox4-mKate2 was generated by replacing TagBFP with mKate2 in the mito-BFP vector (a gift from Prof. Gia Voeltz (Addgene plasmid #49151; http://n2t.net/addgene:49151; RRID: Addgene_49151)). mCherry-Drp1 was a gift from Prof. Gia Voeltz (Addgene plasmid #49152; http://n2t.net/addgene:49152; RRID: Addgene_49152). mCherry2 was first amplified from mCherry2-N1 vector. mKate2 was first amplified from pAAV-CMV-VEGFA-T2A-mKate2 (MiaoLing Plasmid Sharing Platform).

### Sample preparation

To label microtubules in live U2OS cells, we followed a previously described protocol^31^, in which the cells were co-incubated with 4 μM PV-1 and 5μM Tubulin-Atto 488 at 37 °C for 1 hour, then the cells were washed three times with culture medium (warmed to 37°C) and cultured at 37 °C for another 1 hour. Finally, the medium was replaced with phenol red free McCoy’s 5A medium and imaged via DR SPIM.

For labelling microtubules in fixed cells, U2OS cells were seeded onto coverslips at 37 °C with 5% CO_2_ for 16 hours, before fixation, the cells were washed with phosphate buffered saline (PBS, Thermo Fisher Scientific) at 37°C and then treated with fixing buffer (3% Paraformaldehyde (Electron Microscopy Sciences), 0.1% Glutaraldehyde (Electron Microscopy Sciences), 0.2% Triton X-100 (Sigma-Aldrich)) for 15 mins, then incubated with 0.2% Triton X-100 for 15 mins, and blocked with blocking buffer (3% bovine serum albumin (BSA, Sigma-Aldrich) and 0.05% Triton X-100 (Sigma-Aldrich)) for 20 mins at room temperature, after that cells were incubated with an anti-alpha tubulin antibody (Abcam, 1:500 dilution) over night at 4 °C. Subsequently, the primary antibody was removed and the cells were washed twice with PBS. Next, the cells were incubated with a secondary antibody (Abcam, labeled with Alexa Fluor 488, 1:400 dilution) for another 2 hours at room temperature. Finally, the antibody was removed and the cells were washed three times with PBS before imaging.

For labeling the ER in fixed cells, U2OS cells were first transfected with EGFP-Sec61β using Lipofectamine LTX according to the standard protocol and cultured at 37 °C with 5% CO_2_ for an additional 24 h. Before imaging, the cells were fixed with 2% Glutaraldehyde for 20 mins.

For labeling mitochondria in fixed cells, U2OS cells were first transfected with Tomm20-EGFP (mitochondrial outer membrane) or Cox4-EGFP (mitochondrial matrix) using Lipofectamine LTX according to the standard protocol and cultured at 37 °C with 5% CO_2_ for an additional 24 h. Before imaging, the cells were fixed with 2% Glutaraldehyde for 20 mins.

### Quantitative analysis the interactions between the ER and mitochondria

To quantitative analysis the interaction between the ER and mitochondria, we first rendered the volume of the three-dimensional structure of the ER and mitochondria from SR images by Amira. Then, we combined the SR images with the LR images to determine the fusion or fission sites of mitochondria. The y-z slices of the ER and mitochondria at fusion or fission sites were obtained by orthoslice function of Amira software. Finally, we used Fiji^35^ software to analyze the spatial relationships between the ER and mitochondria according to the cross-sectional information of ER-mitochondrial contact sites.

### Image visualization

The three-dimensional time-lapse color code image stacks were generated by using Fiji^37^ plugin and loaded into commercial Imaris software for visualization or directly projection to show the depth information. All the others 3D rendering images were generated by loading the time-lapse image stacks into Amira or Imaris software.

## Notes

### Competing Interest Statement

The authors have declared no competing interest.

## References

1. Phillips, M. J. & Voeltz, G. K. Structure and function of ER membrane contact sites with other organelles. Nat. Rev. Mol. Cell Bio. 17, 69–82 (2015).

2. Eisner, V., Picard, M. & Hajnóczky, G. Mitochondrial dynamics in adaptive and maladaptive cellular stress responses. Nat. Cell Biol. 20, 755–765 (2018).

3. Wong, Y. C., Kim, S., Peng, W. & Krainc, D. Regulation and function of mitochondria-lysosome membrane contact sites in cellular homeostasis. Trends. Cell Biol. 29, 500–513 (2019).

4. Valm, A. M. et al. Applying systems-level spectral imaging and analysis to reveal the organelle interactome. Nature 546, 162–167 (2017).

5. Sahl, S. J., Hell, S. W. & Jakobs, S. Fluorescence nanoscopy in cell biology. Nat. Rev. Mol. Cell Biol. 18, 685–701 (2017).

6. Choquet, D., Sainlos, M. & Sibarita, J. B. Advanced imaging and labelling methods to decipher brain cell organization and function. Nat. Rev. Neurosci, 1–19 (2021).

7. Schermelleh, L. et al. Super-resolution microscopy demystified. Nat. Cell Biol. 21, 72–84 (2019).

8. Huang, B., Wang, W., Bates, M. & Zhuang, X. W. Three-dimensional super-resolution imaging by stochastic optical reconstruction microscopy. Science 319, 810–813 (2008).

9. Huan, F. et al. Ultra-high resolution 3D imaging of whole cells. Cell 166, 1028–1040 (2016).

10. Schermetleh, L. et al. Subdiffraction multicolor imaging of the nuclear periphery with 3D structured illumination microscopy. Science 320, 1332–1336 (2008).

11. Shao, L., Kner, P., Rego, E. H. & Gustafsson, M. G. L. Super-resolution 3D microscopy of live whole cells using structured illumination. Nat. Methods. 8, 1044–1046 (2011).

12. Hein, B., Willig, K. I. & Hell, S. W. Stimulated emission depletion (STED) nanoscopy of a fluorescent protein-labeled organelle inside a living cell. P. Natl. Acad. Sci. USA. 105, 14271–14276 (2008).

13. Bodén, A. et al. Volumetric live cell imaging with three-dimensional parallelized RESOLFT microscopy. Nat. Biotechnol, (2021).

14. Chen, B. C. et al. Lattice light-sheet microscopy: Imaging molecules to embryos at high spatiotemporal resolution. Science 346, 1257998 (2014).

15. Li, D. et al. Extended-resolution structured illumination imaging of endocytic and cytoskeletal dynamics. Science 349, aab3500 (2015).

16. Mahecic, D. et al. Homogeneous multifocal excitation for high-throughput superresolution imaging. Nat. Methods 17, 726–733 (2020).

17. York, A. G. et al. Instant super-resolution imaging in live cells and embryos via analog image processing. Nat. Methods 10, 1122–1126 (2013).

18. Gao, L. et al. Noninvasive imaging beyond the diffraction limit of 3D dynamics in thickly fluorescent specimens. Cell 151, 1370–1385 (2012).

19. Planchon, T. A. et al. Rapid three-dimensional isotropic imaging of living cells using Bessel beam plane illumination. Nat. Methods 8, 417–423 (2011).

20. Chang, B. J. et al. Universal light-sheet generation with field synthesis. Nat. Methods 16, 235–238 (2019).

21. Zhao, T. et al. Multicolor 4D fluorescence microscopy using ultrathin bessel light sheets. Sci. Rep. 6, 26159 (2016).

22. Power, R. M. & Huisken, J. A guide to light-sheet fluorescence microscopy for multiscale imaging. Nat. Methods 14, 360–373 (2017).

23. Weigert, M. et al. Content-aware image restoration: pushing the limits of fluorescence microscopy. Nat. Methods 15, 1090–1097 (2018).

24. Qiao, C. et al. Evaluation and development of deep neural networks for image super-resolution in optical microscopy. Nat. Methods 18, 194–202 (2021).

25. Wu, Y. et al. Three-dimensional virtual refocusing of fluorescence microscopy images using deep learning. Nat Methods 16, 1323–1331 (2019).

26. Fang, L. et al. Deep learning-based point-scanning super-resolution imaging. Nat Methods 18, 406–416 (2021).

27. Wang, Z. et al. Real-time volumetric reconstruction of biological dynamics with light-field microscopy and deep learning. Nat Methods 18, 551–556 (2021).

28. Zhang, H. et al. Exceeding the limits of 3D fluorescence microscopy using a dual-stage-processing network. Optica 7, 1627–1640 (2020).

29. Gustafsson, N. et al. Fast live-cell conventional fluorophore nanoscopy with ImageJ through super-resolution radial fluctuations. Nat. Commun. 7, 12471 (2016).

30. Chen, R. et al. Efficient super-resolution volumetric imaging by radial fluctuation Bayesian analysis light-sheet microscopy. J. Biophotonics 13, e201960242 (2020).

31. Zhang, M. et al. Simple and efficient delivery of cell-impermeable organic fluorescent probes into live cells for live-cell superresolution imaging. Light-Sci. Appl. 8, 73 (2019).

32. Culley, S. et al. Quantitative mapping and minimization of super-resolution optical imaging artifacts. Nat. Methods 15, 263–266 (2018).

33. Friedman, J. R. et al. ER tubules mark sites of mitochondrial division. Science 334, 358–362 (2011).

34. Rowland, A. A. & Voeltz, G. K. Endoplasmic reticulum-mitochondria contacts: function of the junction. Nat. Rev. Mol. Cell Bio. 13, 607–615 (2012).

35. Helle, S. C. J. et al. Mechanical force induces mitochondrial fission. eLife 6, e30292 (2017).

36. Dean, K. M., Roudot, P., Welf, E. S., Danuser, G. & Fiolka, R. Deconvolution-free subcellular imaging with axially swept light sheet microscopy. Biophys. J. 108, 2807–2815 (2015).

37. Schindelin, J. et al. Fiji: an open-source platform for biological-image analysis. Nat. Methods 9, 676–682 (2012).

38. Born, M. & Wolf, E. Principles of optics-electromagnetic theory of propagation, interference and diffraction of light (7. ed.). DBLP, 1999.

