## Supplementary Information for "Multi-color 4D superresolution light-sheet microscopy reveals organelle interactions at isotropic 100-nm resolution and sub-second timescales"

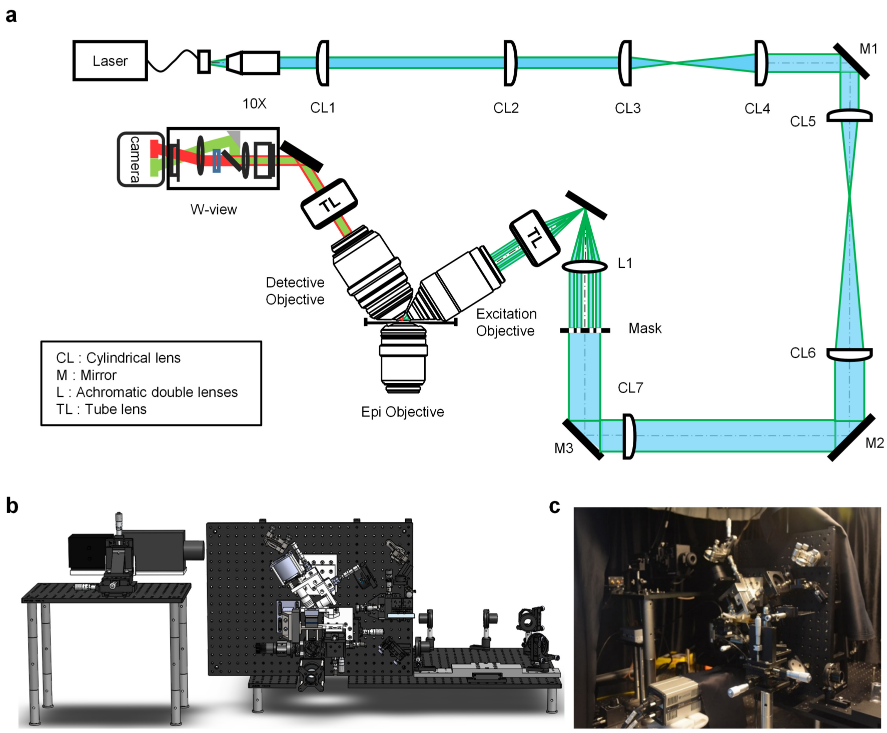


**Supplementary Figure 1 | The dual ring-selective plane illumination microscopy (DR-SPIM) setup. a,** Schematic illustrating the optical path of DR-SPIM. **b,** A 3D solid model of DR-SPIM. **c,** A photograph of DR-SPIM.

| $NA_{outer}$ = 0.33<br>$NA_{inner}$ = 0.18 | Hex 1 | DR-SPIM 1 | Hex 2 | DR-SPIM 2 | Square 1 | DR-SPIM 3 |
|---|---|---|---|---|---|---|
| Pattern at the back focal plane | 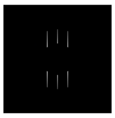 | 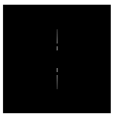 | 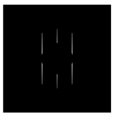 | 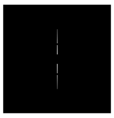 | 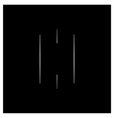 | 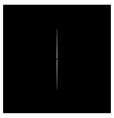 |
| Thickness (µm) | 0.48 | 0.48 | 0.59 | 0.59 | 0.68 | 0.68 |
| Side lobes ratio | 60 % | 35 % | 20 % | 10 % | 3.4 % | 1.3 % |
| Main lobe percent | 44 % | 59 % | 75 % | 85 % | 96 % | 99 % |
| Profile | 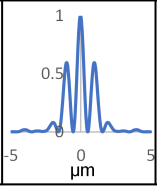  | 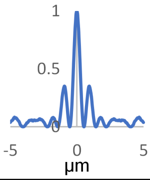  | 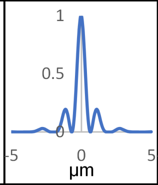  | 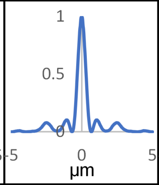  | 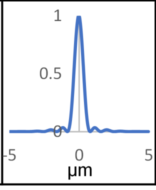  | 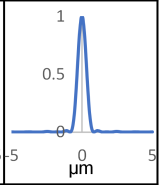  |

**Supplementary Figure 2 | Comparison of the simulated light sheet properties between a dithered lattice light sheet in hex mode (Hex 1 and Hex 2) or square mode[1] (Square 1) and DR-SPIM of the same light sheet thickness (DR-SPIM 1-3).** NA: numerical aperture.

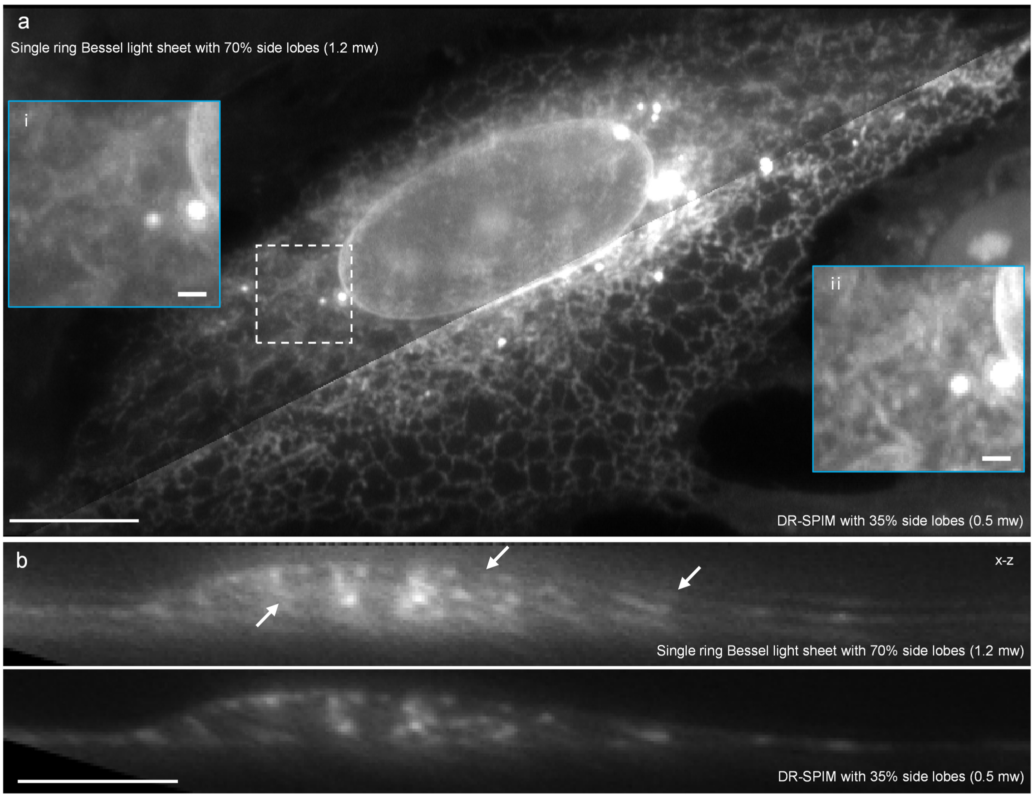


**Supplementary Figure 3 | The endoplasmic reticulum (ER, tagged with EGFP-Sec61β) in a live U2OS cell imaged via single ring Bessel light sheet and DR-SPIM. a,** X-y max intensity projections (MIPs) of the ER imaged via single ring Bessel light sheet[2] with 70% side lobes (left) and DR-SPIM with 35% side lobes (right). Blue boxed regions show the magnified views of the white boxed region imaged via single ring Bessel light sheet (i) and DR-SPIM (ii), respectively. The laser power at the back focal plane was 1.2 mw and 0.5 mw for single ring Bessel light sheet and DR-SPIM, respectively. The exposure time was 10 ms. Scale bar, 10 μm and 1μm (insert). **b,** X-z slices of the ER imaged via single ring Bessel light sheet (top) and DR-SPIM (bottom). The white arrows show artifacts caused by the high side lobes of single ring Bessel light sheet. Scale bar, 10 μm.

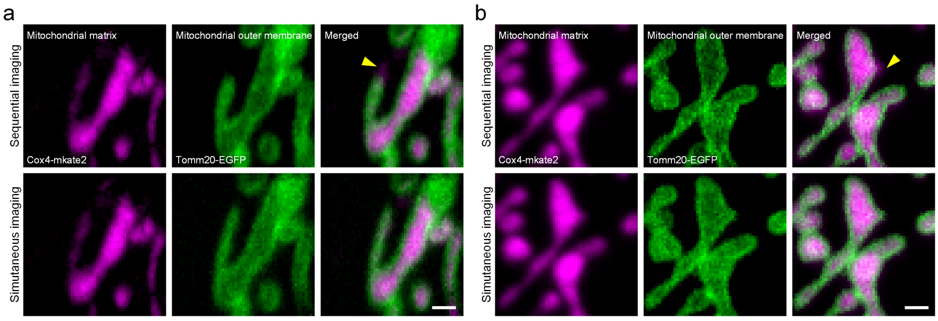


**Supplementary Figure 4 | Comparison of sequential (top rows) and simultaneous (bottom rows) dual-color imaging of mitochondrial matrix (tagged with Cox4-mKate2, red) and mitochondrial outer membrane (tagged with Tomm20-EGFP, green) in a live U2OS cell.** Simultaneous dual-color excitation avoids the misinterpretations (the yellow arrows) caused by the time delay arising from the sequential dual-color imaging. Scale bar, 1 μm.

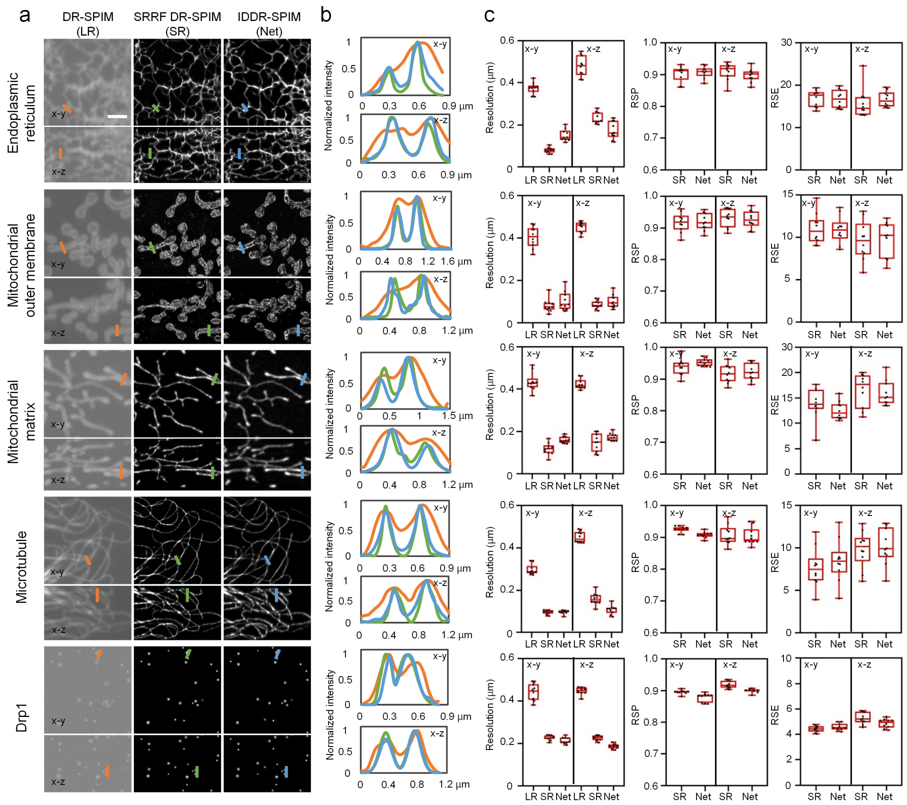


**Supplementary Figure 5 | Comparison of DR-SPIM, SRRF DR-SPIM, and IDDR-SPIM in terms of spatial resolution and fidelity for various organelles and the Drp1 oligomers.** **a,** X-y and x-z MIPs of the indicated organelle or protein imaged via DR-SPIM (LR), SRRF DR-SPIM (SR), and IDDR-SPIM (Net). Scale bar, 2 µm. **b,** Corresponding line profile plots along the lines in **a**. **c,** Corresponding x-y and x-z spatial resolution, resolution-scaled Pearson coefficient (RSP), and resolution-scaled error (RSE)[3] of DR-SPIM (LR), SRRF DR-SPIM (SR), and IDDR-SPIM (Net) reconstructions.

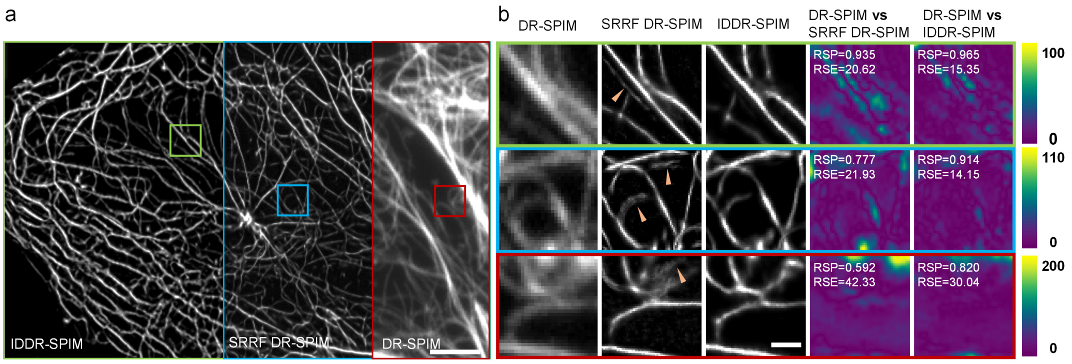


**Supplementary Figure 6 | The rapid motions of the microtubules lead to reconstruction errors of SRRF DR-SPIM. a,** X-y MIPs of microtubules (labelled with Tubulin-Atto 488) in a live U2OS cell imaged via IDDR-SPIM, SRRF DR-SPIM, and DR-SPIM. **b,** Magnified views (left three columns) of the corresponding boxed regions in **a** and their error maps (right two columns) and corresponding RSP and RSE[3] of IDDR-SPIM and SRRF DR-SPIM reconstructions compared to raw DR-SPIM reconstructions. The arrows indicate that the microtubule motion blur leads to the reconstruction errors. The apparent discrepancy between SRRF DR-SPIM and raw DR-SPIM reconstructions is likely to arise from the rapid motions of the microtubules. Scale bar, 1 μm.

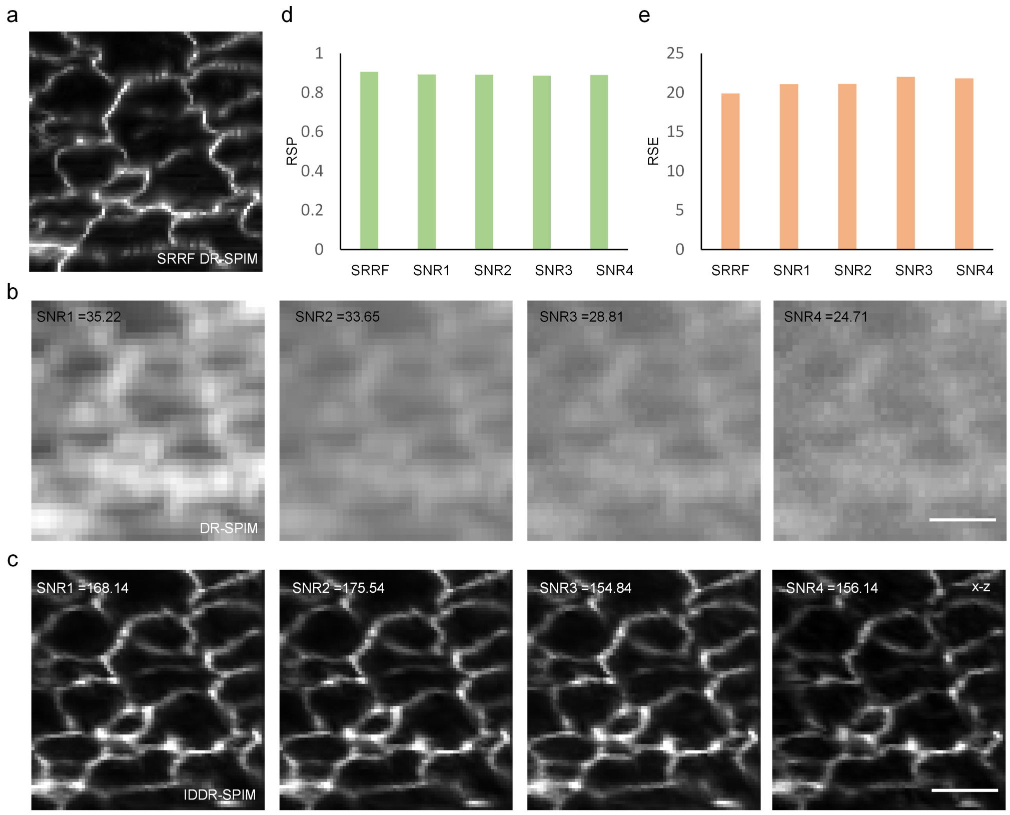


**Supplementary Figure 7 | Reconstruction fidelity of IDDR-SPIM based on the diffraction-limited input images of the ER with different signal-to-noise ratio (SNR). a,** An x-z MIP of SRRF DR-SPIM reconstruction. **b,** The diffraction-limited DR-SPIM reconstructions with different SNR. **c,** The corresponding IDDR-SPIM reconstructions. **d, e,** RSP and RSE of corresponding SRRF DR-SPIM (SRRF) reconstruction in **a** and IDDR-SPIM reconstructions with different SNR (SNR1-4) in **c**. Scale bar, 1μm.

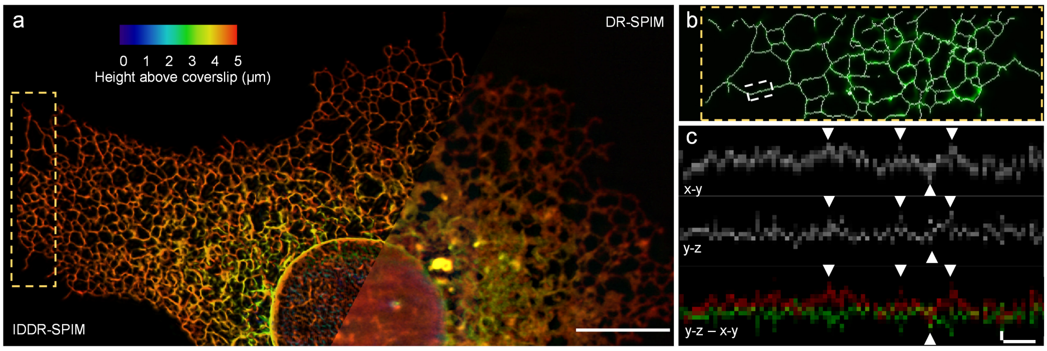


**Supplementary Figure 8 | 3D superresolution imaging of the ER in a live U2OS cell. a,** A color-coded MIP of the ER (tagged with EGFP-Sec61β). The left part shows the superresolution result reconstructed via IDDR-SPIM. The right part shows the diffraction-limited DR-SPIM result. The boxed region was imaged via IDDR-SPIM at a volumetric imaging rate of 17 Hz, and additional frames are shown in **Supplementary Video 5**. Scale bar: 10 μm**. b,** Magnified views of the boxed region in **a**, identified via a skeletonization algorithm. The midline of each ER tubule (white) is mapped onto the fluorescence signal (green). **c,** Positions of the midlines of the white boxed region in **b** in x-y and y-z planes and the merged results of the x-y (red) and inverted y-z planes (green) are plotted as kymographs of oscillation amplitude over time, revealing the oscillations (indicated by the white arrows) of the ER tubules in three dimensions. Scale bar: 100 nm (y axe) and 0.5 s (x axe).

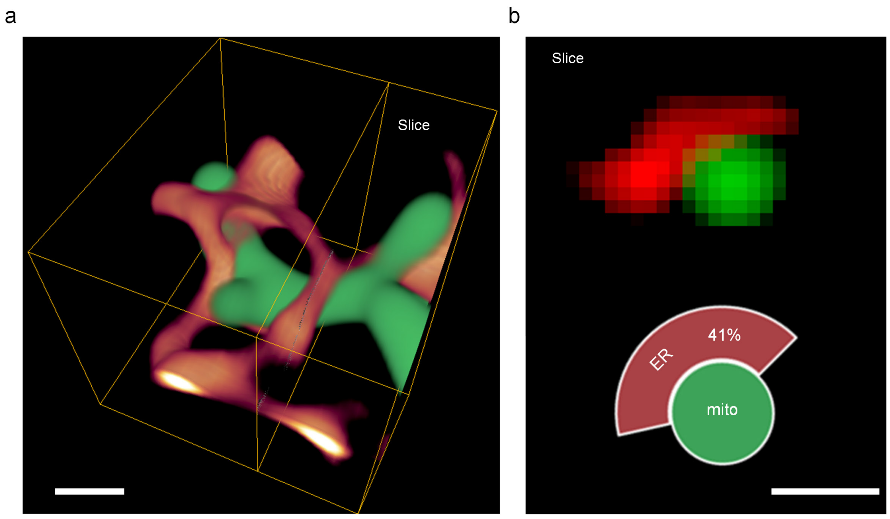


**Supplementary Figure 9 | Quantitative analysis of the interaction between the ER (red) and mitochondria (green). a,** Dual-color 3D rendering shows the interaction between the ER and mitochondria in three dimensions. Scale bar, 500 nm. **b,** A raw reconstructed image (top) of the corresponding y-z sectional plane in **a** and a corresponding 2D model (bottom), showing the percentages of the ER wrapping around the mitochondrial circumference. Scale bar, 500 nm.

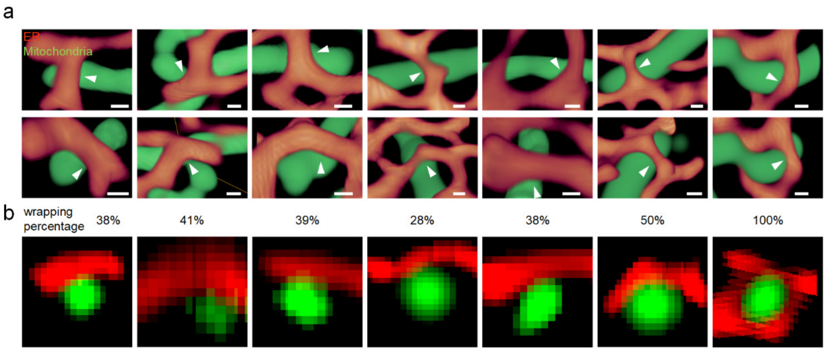


**Supplementary Figure 10 | Mitochondrial morphology at the ER-mitochondria contact sites with similar or different ER wrapping percentages. a,** Dual-color 3D renderings of interactions between the ER (tagged with Sec61β-EGFP, red) and mito Ma (tagged with Cox4-mKate2, green) from two perspectives, respectively, revealing the real spatial relationships and contacts between the ER and mitochondria. The white arrows indicate the ER-mitochondria contact sites. Scale bar, 200 nm. **b,** Raw reconstructed images of the corresponding y-z sectional planes in **a**, showing the percentages of the ER (red) wrapping around the mitochondrial circumference (green).

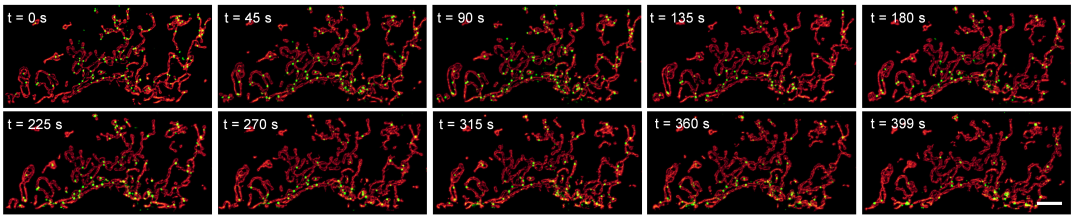


**Supplementary Figure 11 | Representative time-lapse dual-color volume renderings showing the spatial relationships between the Drp1 oligomers (green) and mitochondria (red) imaged via IDDR-SPIM.** Scale bar: 5 μm.

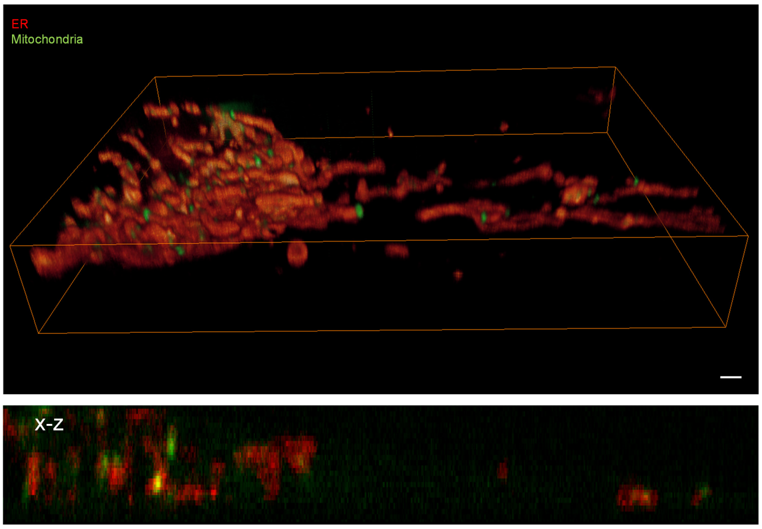


**Supplementary Figure 12 | A dual-color volume rendering (top) and a x-z slice (bottom) of the Drp1 oligomers (green) and mitochondrial outer membrane (red) imaged via spinning disk confocal microscopy (3D mode) in a live U2OS cell.** Scale bar, 2 µm.

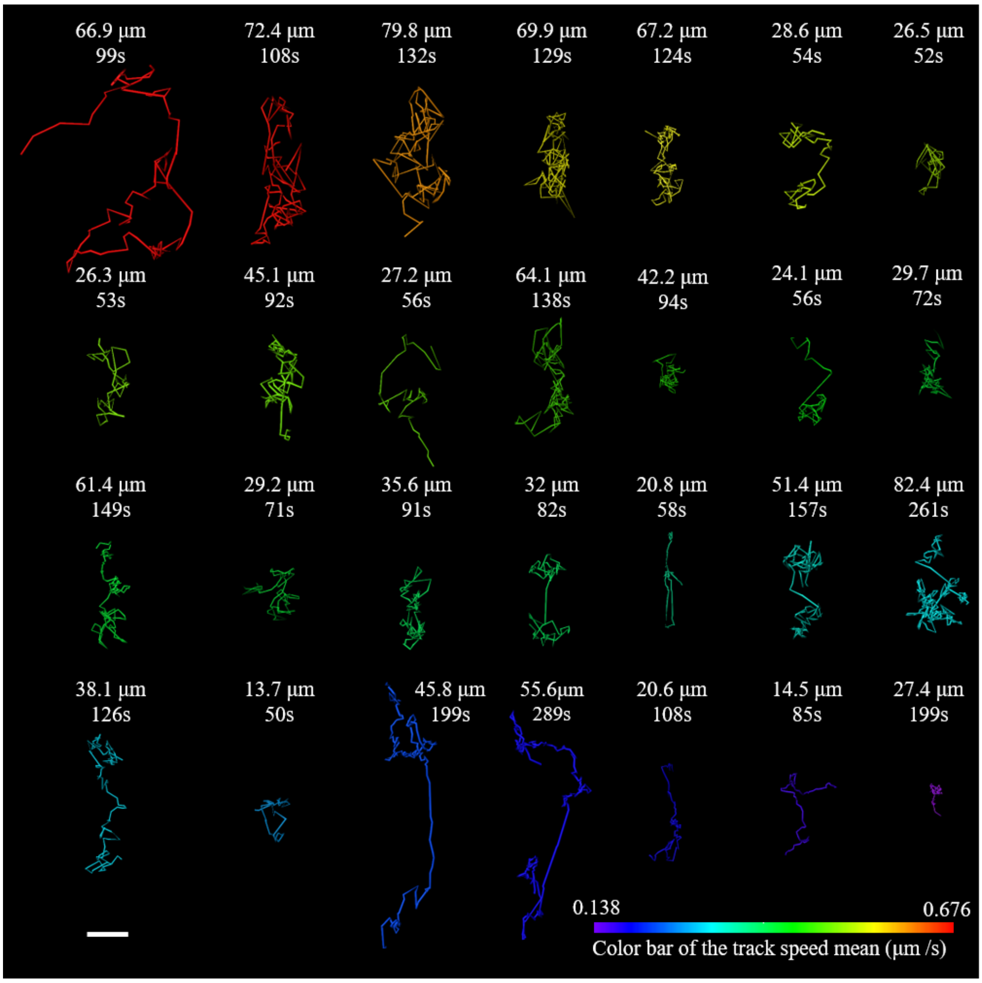


**Supplementary Figure 13 | Trajectories of the Drp1 oligomers not located on the mitochondria.** The numbers indicate the length and tracking time of the corresponding trajectories. Scale bar, 2 μm.

**Supplementary Table 1 | Comparison of the performance between IDDR-SPIM and the other state-of-the-art 3D superresolution methods**

| 3D superresolution microscopy | Lateral resolution (nm) | Axial resolution (nm) | Number of consecutive super resolution images recorded | Illumination intensity | Super resolution frames per second |
|---|---|---|---|---|---|
| 3D-SIM[4] | 110 × 110 | 360 | 40 volumes | ~4-15 W/cm$^2$ | 4 (single color)<br>8.5 (dual color) |
| Instant SIM[5] | 145 × 145 | 350 | ~100 volumes | 5-50 W / cm$^2$ | 34 (single color) |
| Lattice light sheet microscopy in SIM mode[1] | 150 × 230 | 280 | ~340 volumes | ~1-100 μw | ~40 (single color) |
| 3D pRESOLFT[6] | 80 × 80 | 80 | Not available | Not available | 1-2 |
| IDDR-SPIM | 100 × 100 | 100 | 5200 volumes | ~1-100 μw | 200 (single color)<br>400 (dual color) |

### References


1. Chen, B. C. et al. Lattice light-sheet microscopy: Imaging molecules to embryos at high spatiotemporal resolution. *Science* **346,** 1257998 (2014).
2. Zhao, T. et al. Multicolor 4D Fluorescence Microscopy using Ultrathin Bessel Light Sheets. *Sci. Rep.* **6,** 26159 (2016).
3. Culley, S. et al. Quantitative mapping and minimization of super-resolution optical imaging artifacts. *Nat. Methods* **15,** 263-266 (2018).
4. Reto, Fiolka, R., Shao, L., Rego, E. H., Davidson, M. W. & Gustafsson, M. W. M. Time-lapse two-color 3D imaging of live cells with doubled resolution using structured illumination. *P. Natl. Acad. Sci. USA*. **109,** 5311-5315 (2012).
5. York, A. G. et al. Instant super-resolution imaging in live cells and embryos via analog image processing. *Nat. Methods* **10,** 1122-1126 (2013).
6. Bodén, A. et al. Volumetric live cell imaging with three-dimensional parallelized RESOLFT microscopy. *Nat. Biotechnol,* (2021).
